# Crafting Mathematical Models for Type 2 Diabetes Progression: Leveraging Longitudinal Data

**DOI:** 10.1101/2025.02.24.639807

**Authors:** Boya Yang, Arthur S. Sherman

**Affiliations:** Laboratory of Biological Modeling, National Institute of Diabetes, Digestive, and Kidney Diseases, National Institutes of Health, Bethesda, Maryland

**Author notes:** **Address reprint requests to:** Correspondence: Arthur Sherman, 12A South Drive, Room 4007, Bethesda, MD 20892-5621 USA. **Declaration of interest statement:** None.

**Keywords:** Type 2 diabetes progression, Mathematical model, Longitudinal T2D data, Non-linear mixed-effect modeling

## Abstract

Mathematical modeling is a powerful quantitative tool to investigate the pathogenesis of type 2 diabetes (T2D). Most modeling work on the progression of T2D has been formulated by modifying the pioneering model of Topp et al, which established the paradigm of glucotoxicity as the main driver of pathogenesis. However, certain parameter values in the Topp model deviate from physiological data, leading to predictions that deviate from clinical scenarios. Moreover, the simple structure of the model limits its explanatory capacity for clinical data. Leveraging a four-dimensional longitudinal dataset from Southwest Native Americans who progressed from normal glucose tolerance to T2D, we developed a series of models, starting with a minimally modified version of the Topp model and iteratively incorporating additional model elements to account for new biological mechanisms until optimal data fit was achieved. The notable variability of the individual trajectories was overcome by the non-linear mixed-effect modeling approach. Despite the absence of a discernible common trend among the individual trajectories of each variable, the model effectively captured the diverse glucose-insulin dynamics of individuals progressing to T2D. The reliability of the model was reinforced by its successful cross-validation against a subset of individuals progressing only to prediabetes. The systematic model selection process aided in navigating the trade-off between model complexity and practicability, culminating in a robust framework to address controversial questions in the diabetes field in future research.

## 1 Introduction

The pathogenesis of type 2 diabetes (T2D) is complex and involves intricate signaling pathways in insulin secretion and action. Understanding the mechanisms underlying the progression of the disease is essential for developing effective prevention and management strategies to mitigate its impact on healthcare systems and individuals’ quality of life. Nevertheless, due to the intensive requirement for long-term follow-up data collection in humans, very few clinical studies have been conducted to study the evolution of T2D. Consequently, most research in the field has relied on cross-sectional data. Yet, prospective longitudinal studies have demonstrated that individual trajectories exhibited markedly distinct progression profiles when contrasted with data gathered retrospectively and averaged across subjects in cross-sectional analyses [1]. Hence, alternative approaches to studying the personalized development of T2D are desired.

Mathematical modeling has been shown to be an effective quantitative approach to investigating complex biological systems at a relatively low cost. The work of Bergman et al. [2], established a method of defining and estimating the insulin sensitivity index (*S*_*I*_) and a measure of beta-cell function (AIRg) based on an ordinary differential equation (ODE) model depicting the glucose-insulin loop. It has been widely used in clinical research, with over 2500 citations. However, that model only addresses the metabolic state at a moment in time. Other ODE-based models built upon known or hypothesized mechanisms governing the long-term evolution of the glucose-insulin system, are capable of providing a mechanistic understanding of disease progression and the dynamics variations of data variables over an individual’s lifespan. However, only a limited number of such mathematical models have been proposed because of the inherent challenge of capturing the complexity of the disease progression and the paucity of longitudinal data.

The pioneering model investigating the progression of diabetes was formulated by Topp et al [3], adding the slow dynamics of beta-cell mass to the classic glucose-insulin loop to delineate the course of diabetes evolution. De Gaetano et al. [4] employed a similar methodology to construct a model illustrating the concurrent evolution of beta-cell mass, pancreatic beta-cell replication reserve, and insulinemia in pancreatic islet compensation. They then enhanced the model to simulate longitudinal alterations in insulin sensitivity derived from the euglycemic clamp, as well as lifetime fluctuations in glucose and insulin concentrations, and glycated hemoglobin levels during the oral glucose tolerance test [5]. Ha et al. (2016) elaborated the model of Topp et al with further physiological knowledge, delineating the dynamics of beta-cell secretory capacity per cell as an intermediate layer between fast glucose and insulin changes and slower alterations in beta-cell mass. Subsequently, the model was expanded to encompass dynamic glucose flux during meals and OGTTs and insulin granule exocytosis [6] and yielded results consistent with findings from longitudinal studies and offered insights into the pathogenesis of T2D.

Despite the advances in these models, they overlooked the effect of free fatty acids (FFAs) on beta-cell mass and function and insulin sensitivity, simply assuming that sensitivity decreases with time or age. However, numerous studies have supported the hypothesis that excessive FFA load serves as a major disruptor of the glucose-insulin regulatory system, termed lipotoxicity [7, 8], whereas the correlation between insulin sensitivity and age is weak [1]. Yang et al. [9] introduced the impact of obesity into a Topp-like model, which facilitated capturing both the fluctuations and long-term trends of glucose data from Southwest Native Americans (SWNA). Yildirim et al. [10] developed a model, building on the work of Ha et al., that delineated the dynamic interactions between plasma glucose, insulin, free fatty acids (FFA), insulin sensitivity, systemic inflammatory status, beta-cell function, and mass. More recently, De Gaetano et al. [11] proposed a six-dimensional ODE model explicitly considering the contributions of physical exercise and food intake to variations in insulin sensitivity.

When a well-established systems biology model is calibrated with data, it can provide significant assistance in clinical studies and treatment optimization. Due to the intricate pathophysiology of T2D, models delving into the disease progression are typically complex and necessitate calibration using individualized longitudinal data to ensure their applicability in investigating individualized disease progression. Yet, none of the models in the literature have been calibrated using individualized longitudinal time-series data exceeding two dimensions, whereas a high data dimension is required for the calibration of complex models to prevent overfitting.

To tackle these problems, we leveraged a four-dimensional longitudinal T2D dataset from SWNA and developed a series of obesity-diabetes models incorporating different biological mechanisms and assumptions regarding the glucose-insulin regulatory system. The optimal model was determined by comparing models’ performance in characterizing markedly distinct individual progression profiles. Despite the absence of a discernible common trend among the individual trajectories of each variable, the model effectively captures the diverse glucose-insulin dynamics during T2D development. The credibility of the model is reinforced by its successful validation against a subset of individuals progressing only to prediabetes. This robust modeling framework enables the testing of hypotheses about the underlying causes of the disease. We provided one example of how the models can be employed to address controversial questions in the diabetes field.

## 2 Study design

This study is a secondary analysis of a subset of participants drawn from a longitudinal epidemiological investigation carried out in a Southwest Native American (SWNA) community in Arizona, USA [12]. This conducted from 1965 to 2007, involved individuals aged 5 years and above, who were periodically invited for outpatient research assessments approximately every two years. These assessments included an oral Glucose Tolerance, along with evaluations of body mass index (BMI) and percentage of body fat (PFAT). As previously described [13], adult community members without T2D were recruited. These individuals were screened for general health and medication review one month before each assessment, ensuring the exclusion of medications known to influence glucose or insulin metabolism.

The subgroup of the participants who had at least 3 OGTTs was previously analyzed by Ha et al for a different purpose [14]. The data from these 201 individuals will be referred to as “the complete dataset”. To analyze the progression of T2D, we selected individuals from the complete dataset who had both normal glucose tolerance and normal fasting glucose (NGT/NFT) at baseline and developed prediabetes or diabetes during the following visits (Table 1). The selection of individuals with prediabetes and diabetes is based on threshold crossings of 2-h glucose, as the participants in the study had a low incidence of impaired fasting glucose with normal glucose tolerance compared to impaired glucose tolerance with normal fasting glucose, as previously reported for this population [15].

**Table 1.** Baseline characteristics of the datasets.

| Characteristics | Age | Sex<br>Male<br>(%) | PFAT | BMI<br>(kg/m<sup>2</sup>) | FPG<br>(mg/dl) | FPI<br>(μU/ml) | Matsuda<br>Index | $\frac{AUC_I}{AUC_G}$ | Follow-up<br>(years) |
| --- | --- | --- | --- | --- | --- | --- | --- | --- | --- |
| T2D dataset<br>N=17 | 24.8<br>(6.6) | 35.3 | 36<br>(7.8) | 34<br>(6.9) | 88.4<br>(10.1) | 37.4<br>(19.8) | 1.2<br>(0.6) | 1.3<br>(0.8) | 6<br>(2.7) |
| PreDM dataset<br>N=63 | 24.8<br>(5.9) | 61.9 | 31.6<br>(8.6) | 33.8<br>(6.6) | 91<br>(8.7) | 33<br>(18.3) | 1.4<br>(0.8) | 1.2<br>(0.7) | 5.6<br>(2.5) |

The prediabetes progression dataset comprises 63 individuals who were euglycemic (FPG < 100 mg/dl [5.6 mmol/L], 2h-PG < 140 mg/dl [7.8 mmol/L]) at baseline and developed prediabetes (2h-PG ≥ 140 mg/dl) during a follow-up period of 5.6±2.5 years. The T2D progression dataset includes 18 individuals who transitioned from euglycemia to diabetes (2h-PG ≥ 200 mg/dl) during 6±2.7 years of follow-up. One individual was excluded from this dataset due to incomplete data.

This study proposes 20 ordinary differential equation (ODE) models with three or four dependent dynamic variables and incorporating different combinations of eight biological mechanisms. The T2D progression dataset was employed to select the optimal model for the investigation of disease development, while the prediabetes progression dataset was used for cross-validation of the optimal model. We used the Bayesian Information Criterion (BIC) [16] as a key metric for model selection and evaluation, with the aim of imposing a heavy penalty on model complexity. BIC is defined as follows:

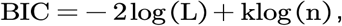

where L denotes the maximum value of the likelihood function of the model; k represents the number of parameters; and n is the sample size.

The data variables used in the models are age, percent body fat (PFAT), fasting plasma glucose (FPG), fasting plasma insulin (FPI), the Matsuda index, and a form of the insulinogenic index (IGI) defined by AUC_I_/AUC_G_ during the OGTT. Age and PFAT were regarded as time-varying parameters, the values of which were imported into the model directly from the data. Model calibration was performed using the four-dimensional longitudinal data of each individual. We observed a great deal of variability of individual trajectories and employed the non-linear mixed-effect (NLME) modeling approach to mitigate this.

PFAT was chosen over BMI as an obesity indicator, since BMI tends to decrease as diabetes progresses, whereas PFAT remains unaffected [17]. The ratio AUC_I_/AUC_G_ stands for the rate of the area under the curve of insulin to the area under the curve of glucose during the three-hour OGTT and serves as an index for evaluating beta-cell function. The Matsuda index, an indicator of insulin sensitivity, was calculated with the formula [18]:

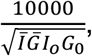

where G_0_ and I_0_ denote the fasting glucose (mg/dl) and fasting insulin (µU/ml), and 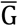 and Īstand for the average glucose and insulin over the OGTT calculated as AUC/120 min by the trapezoidal rule. To limit the number of estimated parameters, we focused on delineating the long-term dynamics of metabolic variables at their basal states, while omitting the postprandial state dynamics. Other than calculating the Matsuda index and insulinogenic index, only the fasting glucose and insulin data were fitted to the model.

## 3. Model selection and results

Our approach for constructing the mathematical models is to begin with a simplest conceptual model and then progressively introduce complexity until the model can account for data well. The model of Topp et al employs the simplest structure to illustrate the main features of the glucose-insulin regulation system: fast negative feedback by insulin to control glucose on a time scale of hours, slow negative feedback by beta-cell mass to compensate for reduced insulin sensitivity, and progression to diabetes when the slow negative feedback tips over into positive feedback. The three differential equations and auxiliary functions are:

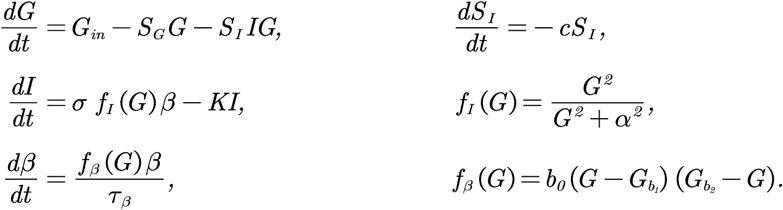

In the model, G (mg/dl), I (μU/ml), and β (mg) denote the plasma glucose concentration, insulin concentration, and the mass of functional beta-cells (active for appropriate insulin production and secretion) at time t (days), respectively. The parameter G_in_ represents the average rate of glucose influx per day. The term S_G_G denotes insulin-independent glucose uptake, while S_I_IG represents insulin-dependent glucose uptake. The coefficient S_I_ (ml/μU/day) represents insulin sensitivity, following the concept introduced with the minimal model of Bergman, Cobelli, and colleagues[19]. Insulin secretion from beta-cells is assumed to be stimulated by elevated glucose levels, and the coefficient represents the secretory capacity per beta-cell. The rate of insulin clearance is denoted by k (per day). The functional beta-cell mass is determined by an exponential growth equation with a glucose-dependent growth rate. That rate is postulated to be a downward parabolic function. The product of and represents the total capability of beta-cells to respond to glucose stimulation. To align mathematical expressions with clinical usage, we designate σ β as beta-cell function and σ as beta-cell secretory capacity per cell in this context.

The simplified character of the model of Topp et al restricts its ability to represent clinical observations or experimental results of diabetes [9, 20]. Enhancements can be made by calibrating parameter values using empirical data, replacing specific model elements, and integrating new components that depict additional biological mechanisms, as we detailed in Section 6.1. With minimal adjustments to Topp’s model, we established Model 1. To further enhance model performance, we proposed nine new formulations for the eight model components of Model 1 to enhance the model performance (see Section 6.2 for further details). These new formulations can be viewed as additional model ingredients to augment the base model. Our next step involves selecting an optimal combination of these model ingredients, striking a balance between performance and complexity. The significance of each model ingredient varies and can be assessed by comparing the model BIC values before and after integrating the specific ingredient. However, it is notable that the significance of model ingredients is not strictly additive, as there may be instances of agonistic or antagonistic effects when they are combined.

We started with incrementally adding the ingredients to Model 1 (see Section 6.1.2 for model formulation) and recorded the corresponding BIC values for each iteration (Figure S2). Once the ingredient resulting in the lowest BIC values was identified, we designated the resulting model as Model 2, while all prior trial models were categorized as “the first-generation model”. Subsequently, we systematically incorporated each non-selected ingredient or combinations thereof into Model 2 until achieving a model with a substantial decrease in BIC. This refined model was designated as Model 3, and trial models established upon Model 2 were labeled as second-generation models. This selection process is schematically illustrated in Figure 1. We selected the model parameters for estimation based on their influence on the model output and the prior information we obtained for these parameters. The impact of parameters on the outcome was inferred from the results of the extended Fourier Amplitude Sensitivity Test (eFAST) method [21], as shown in Figure S6.

**Figure 1.**
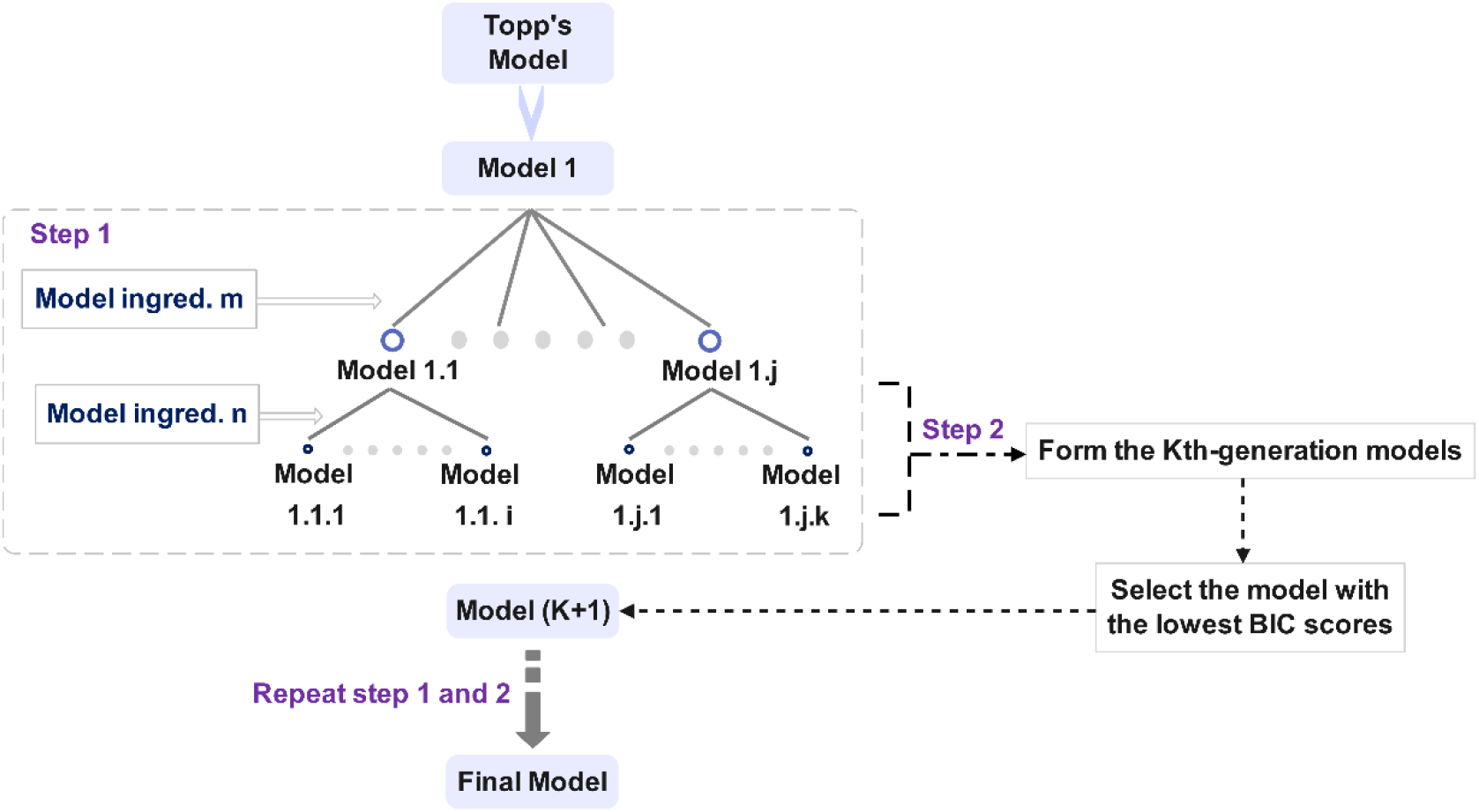
Schematic illustration of the model selection procedure.

To highlight the primary ingredients refining model performance, we ranked the sets of model ingredients based on their impact on reducing the BIC values. The sequence of decreasing BIC values exhibited a trend toward convergence after reaching the fifth generation (Figure 2). In contrast to the 46.5% decrease in BIC observed between the first- and second-generation models, the BIC values of the fifth-generation models exhibit a maximum variation of 2.2%, with Model 5 and Model 5.2 ranking the two lowest (Figure S2). The observation vs. prediction plots generated by Model 5 (Figure 3) illustrate its adeptness in capturing the long-term trends of each variable, despite the high inter-individual variabilities in the four-dimensional data. This implies that the model effectively captures the essential features of the glucose-insulin regulatory system. The disparity between Model 5 and Model 5.2 lies in their respective assumptions regarding the impact of excess FFAs on beta-cell secretory capacity (σ), either as solely lipotoxic or exhibiting both positive and negative (biphasic) effects. Despite the marginally lower BIC value of Model 5.2 compared to Model 5, we chose Model 5 for further analysis based on prior clinical evidence supporting the biphasic-effect hypothesis [22]. Therefore, Model 5 was deemed the optimal model for this study.

**Figure 2.**
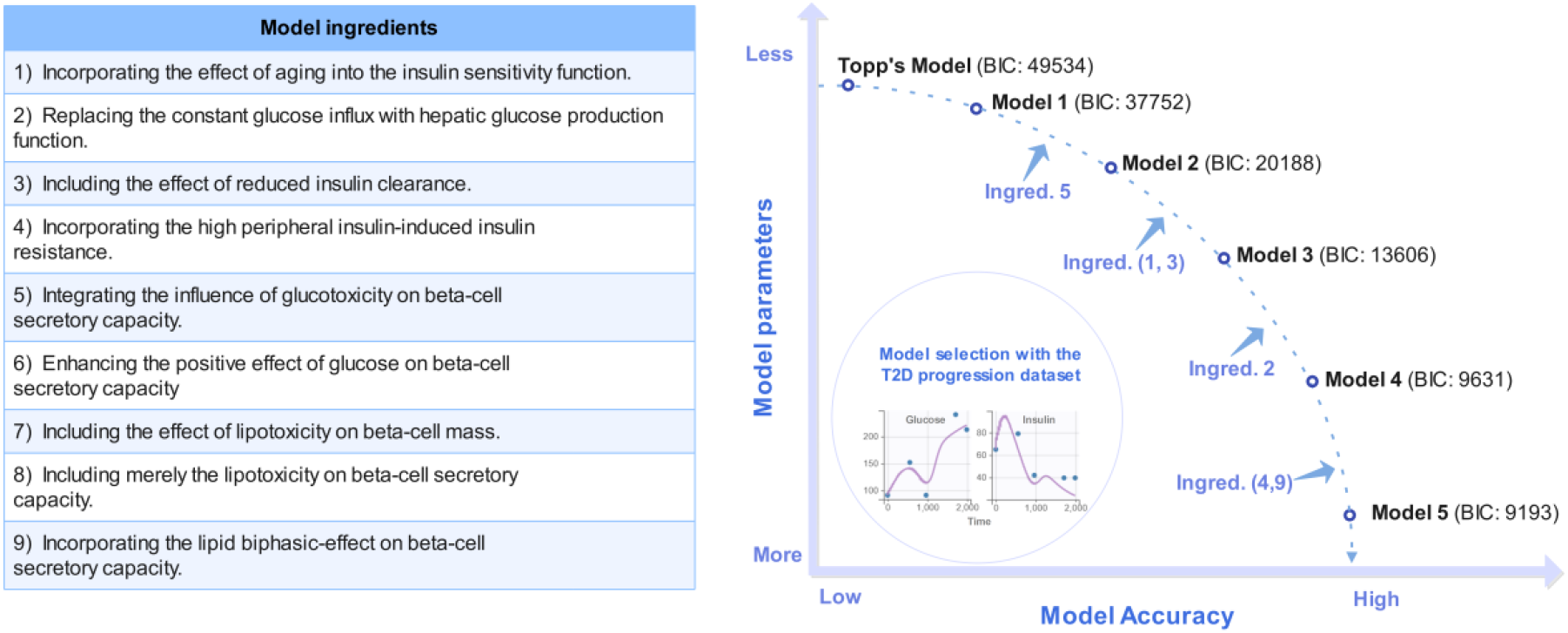
Formulation development for the optimal T2D progression model. Model 1 requires further refinement to effectively elucidate the longitudinal data of SWNA. The left table outlines nine model ingredients that can be added to Model 1 to account for additional biological mechanisms. To strike a balance between model complexity and accuracy, 19 models were derived from Model 1 for performance comparison, each incorporating different subsets of these model ingredients (Figure S2). The decreasing trend in BIC values indicates a convergence toward optimal model performance by the fifth generation of models.

**Figure 3.**
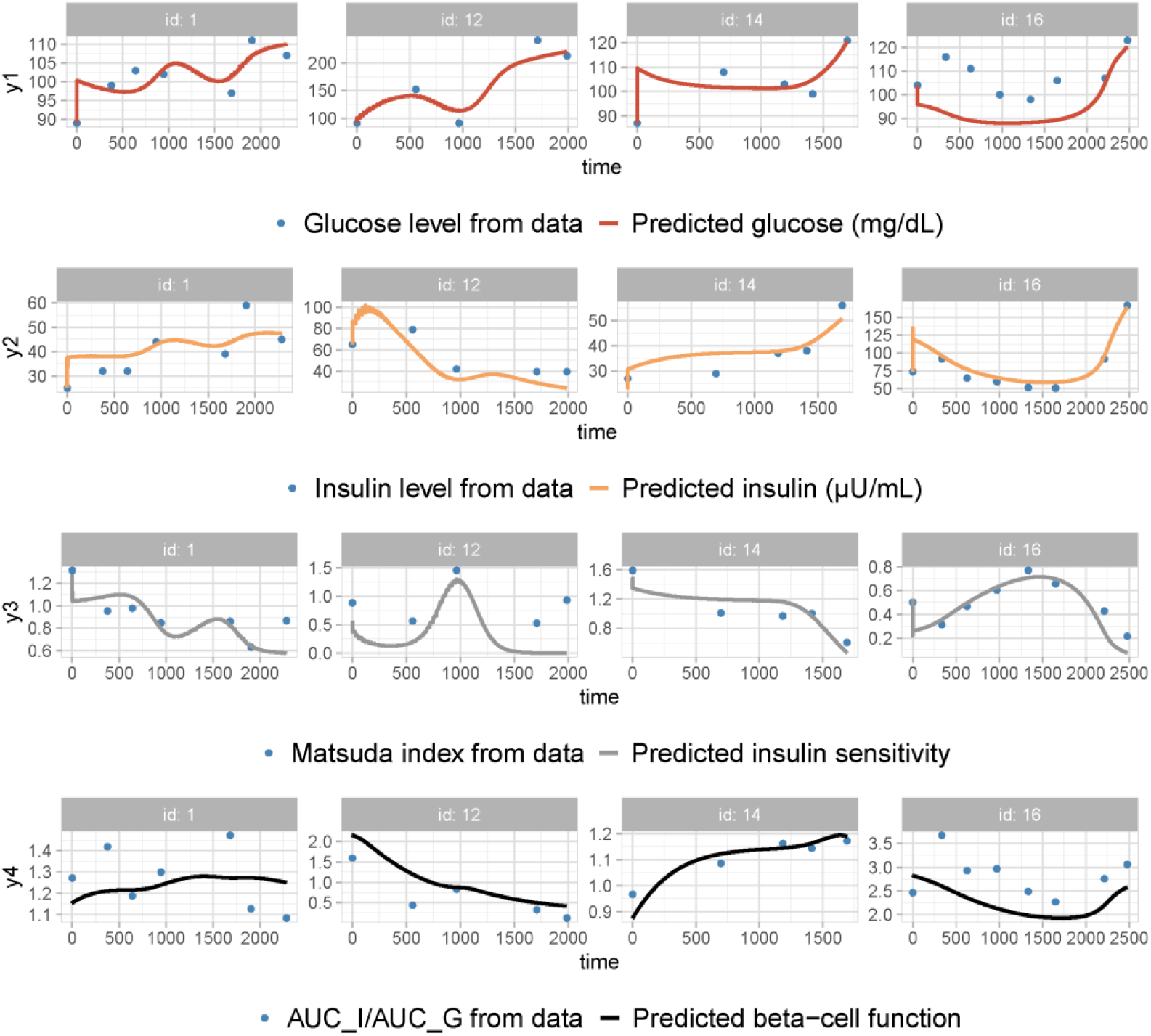
Observation vs prediction plots generated by applying Model 5 to the T2D progression dataset. The time unit is in days. Plots for Patient ID 1, 12, 14, and 16 are shown as representatives of the entire cohort (for the complete fitting results, refer to the Model 5 report file in the supplemental materials). The fitted variables are fasting glucose *G*, fasting insulin *I*, insulin sensitivity *S*_*I*_, and beta-cell function *σβ*. The time is measured in days. The data are shown as blue dots, and the solid curves represent the predictive dynamics for each variable. Despite the distinct data excursion profiles for each patient, the model predictions effectively capture the trend of variations for all data variables.

Considering the relatively small size of the T2D progression dataset, we further validated Model 5 using the prediabetes progression dataset through a three-fold cross-validation. K-fold cross-validation is a commonly used approach for testing models’ predictive ability while flagging problems like overfitting or selection bias. However, this approach is seldom applied to NLME models for performance evaluation, primarily due to potential disparities in the distribution of individual parameters across the three-fold subsets. [23]. To address this challenge, we applied the population parameter distributions obtained from the training dataset to the test dataset, estimating only the random effects (individual variations from the population) on the test dataset (Figure 4). Subsequently, the BIC values derived from this approach on the test dataset (labeled as “BIC by testing”) were compared with the BIC values obtained by fitting the test dataset with both fixed effects and random effects (labeled as “BIC by fitting”). This process was repeated three times on the three test datasets.

**Figure 4.**
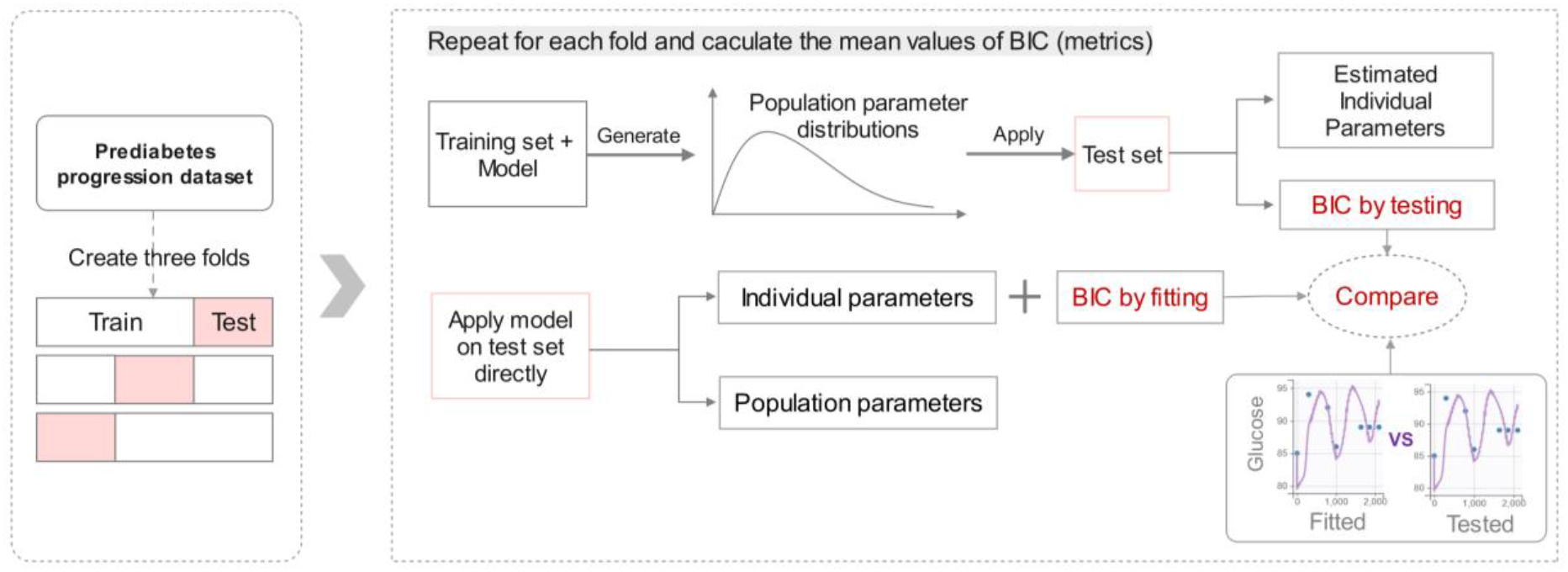
Illustration of the approach used for evaluating NLME model performance through three-fold cross-validation. The prediabetes dataset was evenly divided into three test datasets. The complementary sets of each test dataset were labeled as training datasets. BIC values obtained by fitting and testing with the three test datasets were then averaged for comparison.

The rationale behind this approach is that if the BIC by testing closely aligns with the BIC by fitting for a model exhibiting good performance, it suggests this favorable model behavior is achieved without overfitting, as the number of estimated parameters is drastically reduced without estimating fixed effects. The observation vs prediction plots on the prediabetes progression training dataset (Figure 5) demonstrate that Model 5 accurately characterizes the progression patterns of the metabolic state for each individual. By comparing the BIC values obtained by fitting and testing with the three test datasets (Table S2), we can conclude that the favorable model performance is free from concerns regarding over-parameterization.

**Figure 5.**
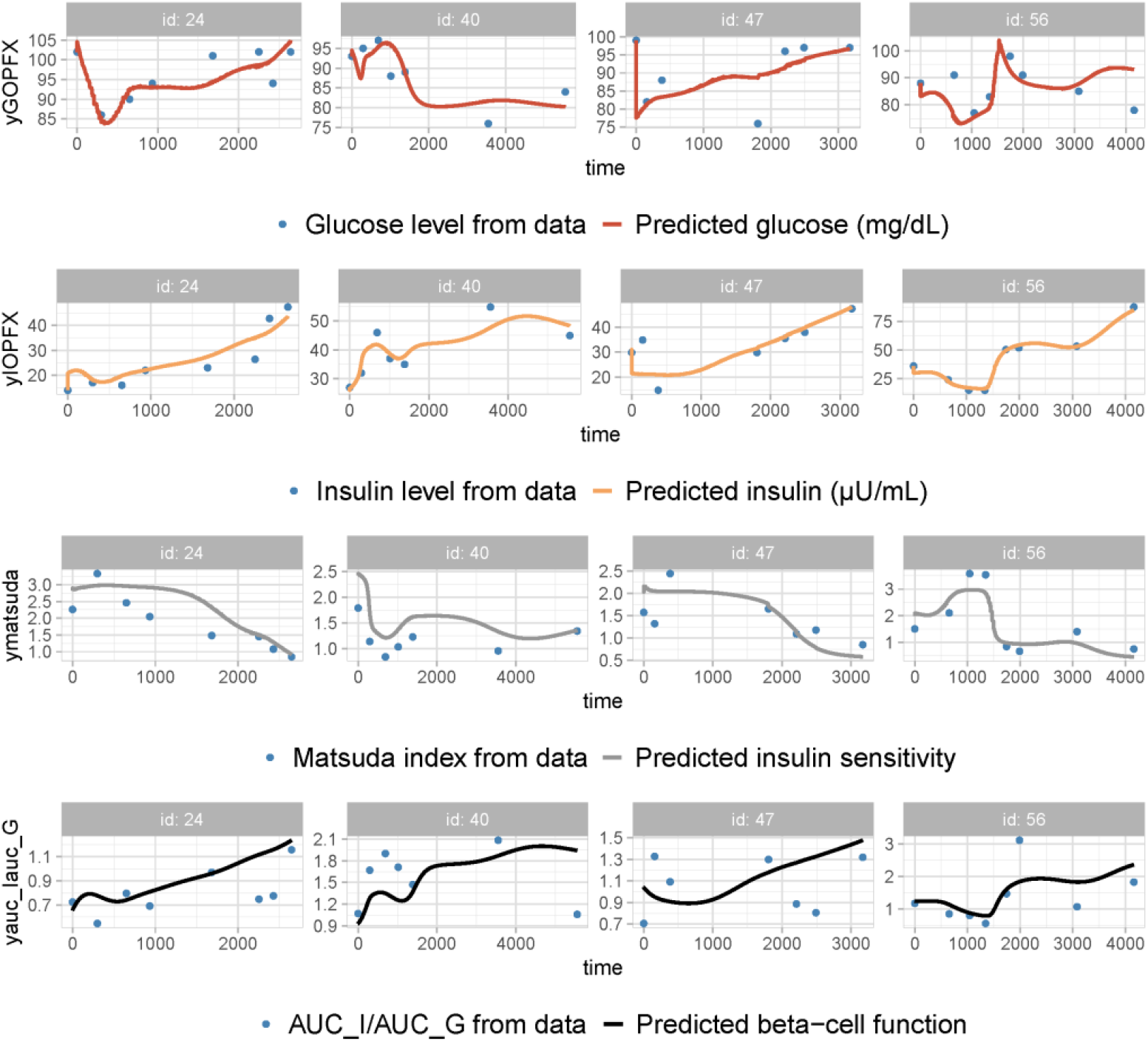
Observation vs prediction plots generated by applying Model 5 to the prediabetes progression training dataset 1. Plots for Patient ID 24, 40, 47, and 56 are shown as representatives of the entire cohort (for the complete fitting results, refer to the report files for cross-sectional validation in the supplemental material). The fitted variables are fasting glucose *G*, fasting insulin *I*, insulin sensitivity *S*_*I*_, and beta-cell function *σβ*. The time is measured in days. The data are shown as blue dots, and the solid curves represent the predictive dynamics for each variable.

## 4 Prospect of model implications

The strong model performance on the chaotic dataset for both prediabetes and diabetes progression suggests that Model 5 effectively captures the glucose-insulin regulatory system across all disease states. Thus, the biological mechanisms incorporated into Model 5 are essential for analyzing the pathogenesis for the SWNA population. In particular, when comparing estimated individual parameter distributions with the dataset on prediabetes progression and diabetes progression, we found that only the parameter values for glucotoxicity (σ_6_) are significantly different between the two groups (Figure S3). This demonstrates that impaired beta-cell function, in this population that has widespread severe insulin resistance, is the major factor driving the evolution of T2D.

The pathogenesis of T2D has long been debated. The prevailing perspective suggests that, in the presence of insulin resistance, beta-cell dysfunction occurring early in the disease progression is the critical abnormality. An alternative hypothesis is that primary beta-cell overstimulation results in insulin hypersecretion, subsequently leading to insulin resistance and, ultimately, beta-cell failure [24]. Advocates of the insulin hypersecretion theory point to empirical evidence that 1) in the prediabetic state, the insulin concentration during glucose tolerance tests can still be twice as high as in euglycemic lean individuals. 2) fasting glucose does not increase much during the prediabetic phase when basal insulin levels rise dramatically [25, 26]. Therefore, proponents postulate that insulin hypersecretion leads to high glucose rather than the reverse.

However, arguments 1 and 2 above only demonstrate correlation, not causation. Here, we employ our simplest model (Model 1) to quantitatively illustrate why insulin hypersecretion can be driven by increases in glucose that are too small to measure.

We first show that the rapid decrease in insulin sensitivity does cause a large and rapid increase in glucose. Figure 7 (a) and (b) describe the changes in basal glucose and insulin when insulin sensitivity is changed in a step-wise fashion, decreasing by 30% at day 10 and restored to baseline at day 12 (Figure S4(d)). Both insulin and glucose rise significantly as beta-cell mass cannot promptly compensate for insulin resistance (Figure S4(c)).

**Figure 7.**
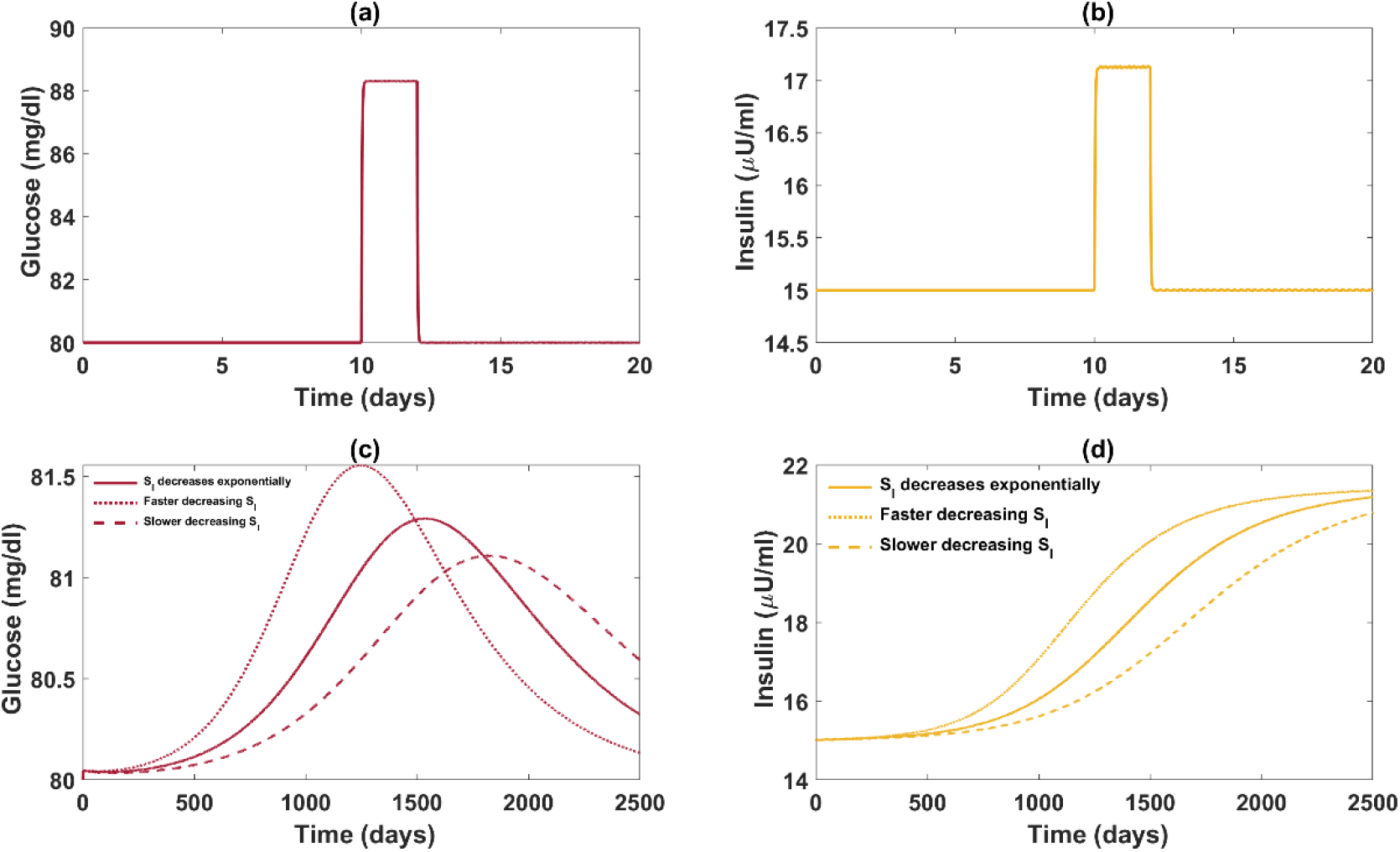
Comparison of glucose-insulin dynamics between fast and slow declines in insulin sensitivity. Refer to Figures S4 and S5 for the complete *GI*β dynamics and the plot of insulin sensitivity functions. Panels (a) and (b) describe the changes in basal glucose and insulin when insulin sensitivity is formulated as a step function with a 30% decrease during day 10 and day 12. Panels (c) and (d) depict the variations of basal glucose and insulin when insulin sensitivity decreases exponentially over the years to the same extent as in panels (a) and (b). The dotted and dashed curves depict the glucose and insulin dynamics associated with higher and lower rates of decrease in insulin sensitivity respectively, compared to the solid curve.

In contrast, when insulin sensitivity gradually decreases over the years, as in the natural history of T2D, the elevation in glucose is insignificant (Figure 7(c)) and diminishes over time (Figure S5(a)), while there still is a 42.3% increase in basal insulin. The rise in glucose is prevented as beta-cells have sufficient time to compensate for insulin resistance (Figure S5(c)). The slower the fall of insulin sensitivity, the smaller the rise in glucose. Similar results can be obtained by the even simpler original model of Topp et al without the additional assumptions and parameter changes made here. The mathematical model, grounded in physiology, reveals that even very slight elevations in glucose can lead to substantial increases in insulin when the slow negative feedback of the regulatory system is activated.

Additionally, we investigate how varying rates of decline in insulin sensitivity can affect the glucose regulatory system. It is noteworthy that a robust beta-cell response does not preclude the possible presence of beta-cell dysfunction, interpreted here as the inability of beta-cells to counteract insulin resistance effectively to maintain glucose homeostasis. Figure 7(d) shows that a faster decrease in insulin sensitivity can lead to a stronger compensation in insulin secretion between the endpoints. However, the product of insulin and insulin sensitivity (the disposition index) decreases as insulin sensitivity declines faster (Figure S5(e)). This indicates that an enhanced beta-cell response can coexist with a reduced ability to attain glucose homeostasis. Thus, beta-cell hyperresponsiveness can coexist with beta-cell dysfunction. In conclusion, beta-cell hyperresponsiveness in the absence of a noticeable change in glucose before the prediabetic stage does not substantiate arguments for the insulin hypersecretion hypothesis.

Furthermore, Figure 7 (c) and (d) also illustrate the limitations of studying disease progression from cross-sectional data. As individuals progress at different rates, data collected at a specific time point for all subjects may yield misleading information. For instance, if Figure 7 (c) depicts glucose trajectories for three subjects, and data were only collected on day 2500, we might conclude that the subject with slower development in insulin resistance has higher glucose but would draw the opposite conclusion if the data was collected on day 500.

## 5 Discussion

Longitudinal studies investigating the development of obesity and T2D have revealed that the signature variables associated with diabetes progression, such as fasting insulin levels, vary significantly across subjects, exhibiting different initiation points, rates of progression, and magnitudes of changes in trajectories [1]. Hence, investigations based on relevant cross-sectional or average data across subjects may be inadequate to reveal the diverse pathways toward T2D. Moreover, the high within-subject variability requires the collection of frequently sampled, long-term follow-up data, posing a formidable challenge for clinical studies. However, this challenge can be effectively addressed using mathematical models established upon the biological mechanisms of the glucose regulatory systems. T2D progression models calibrated with individual data can provide complete trajectories of each variable over a subject’s lifespan, bridge the gaps between data samples, and enable the comprehensive analysis of personalized disease progression.

Most models studying or involving the evolution of diabetes have been built upon the model of Topp et all, which describes the long-term regulation of glucose and insulin with the simplest structure. These models retained certain aspects of Topp’s model while introducing new elements to address specific questions of interest. Some of the models directly adopt the parameter values in the Topp model for convenience. However, we found that the parameter values used in Topp’s model do not align well with human experimental data. In this work, we selected the model functions and parameter values by leveraging experimental results and clinical data. We elaborated the formulations of our model components, along with corresponding references, to facilitate the development of future modeling studies for T2D.

Moreover, the overly simple model of Topp et al, while effective for illustrating novel concepts, has a limited ability to capture clinical data. Numerous studies have consistently highlighted the substantial impact of excess FFAs on the glucose-insulin regulation system. Therefore, a model confined to only the variables of glucose and insulin overlooks a crucial regulator of the system. Clinical findings in recent decades have provided detailed information for reference in model formulations including high insulin-induced insulin resistance, reduced insulin clearance, the coexistence of positive and negative effects of FFAs on beta-cell function, and so on. On the other hand, incorporating these features into a model can greatly increase the complexity of the model structure. To maximize model efficacy in clinical applications, modelers must strike a delicate balance between model complexity and practicability, a task that requires thorough calibration with sufficient data.

Utilizing a four-dimensional longitudinal SWNA data set, we developed a series of models, commencing with a simple model and iteratively incorporating additional model elements to account for new mechanisms until achieve optimal data fit. As various combinations of model ingredients yield different performance outcomes, we considered twenty different models for comparison, and the key model elements substantially lowering model BIC values were identified. This systematic model selection process informs modelers about the indispensable elements that should be integrated into their models to ensure practicability.

Specifically, upon analyzing the data, we observed that Matsuda insulin sensitivity index correlates better with the percentage of body fat than with age, although we found that adding age to the percentage of body fat further improved the model. This underscores the importance of incorporating the effect of excess FFAs (parameter X) into the insulin sensitivity function within the model. Due to a lack of data, prevailing formulations of the insulin sensitivity function in the field often take the form of an exponential decay function over time. Such an approach does not readily allow incorporating individual variations in time of adiposity and hinders the ability of models to explain data and generate accurate predictions. We proposed two new formulations for the insulin sensitivity function, one is a decreasing function dependent solely on X, while the other exhibits a decline with both X and age. Having the insulin sensitivity function dependent on X considerably reduced the BIC values, and the performance was further enhanced with S_I_(X,AGE).

Another significant advancement in the glucose equation can be achieved by substituting a constant hepatic glucose production (HGP) rate with a dynamic HGP function. A few T2D progression models use a constant basal hepatic glucose production rate for simplicity. Nevertheless, clinical data have demonstrated that basal HGP is overtly elevated concurrently with the upsurge of glucose at the onset of diabetes [17, 27, 28]. To capture the evolution of basal HGP, we formulated the rate of HGP as a function dependent on the level of insulin and excess FFAs. This inclusion links peripheral insulin resistance with hepatic insulin resistance, and the generated HGP-insulin dose-response curve (Fig. 11) aligns well with experimental observations [29, 30].

In many T2D progression models, the insulin clearance rate is assumed to remain constant. However, clinical findings have revealed a rapid reduction in insulin clearance as insulin secretion increases in insulin-resistance states [31]. Considering the data on insulin clearance rate [32], we adopted a simple hyperbolic function dependent on insulin concentration. This choice is bolstered by substantially reduced BIC values. Hence, we replaced the constant insulin clearance rate with this rate function.

The greatest enhancement of model performance the model was achieved by incorporating the impact of glucotoxicity on the dynamic beta-cell secretory capacity σ. Studies have indicated the slow compensation for insulin resistance cannot be solely attributed to mass expansion [20]. Thus, it is necessary to include a dynamic equation for σ in the model, with a rate of change situated between fast changes in glucose and insulin, and slower alterations in beta-cell mass. Upon comparing estimated individual parameter distributions with datasets on prediabetes progression and diabetes progression, we observed significant differences only in the parameter values for glucotoxicity between the two groups. We interpret this as indicating that decreased insulin sensitivity alone does not lead to diabetes unless the beta-cell function is also impaired. Conversely, typical variation in beta-cell function does not generally lead to diabetes in the absence of insulin resistance. The varied disease progression rates among individuals primarily stem from different beta-cell susceptibility to elevated glucose levels.

After integrating the four crucial model elements mentioned above, we advanced to the fourth-generation models, which achieved further refinement of performance through the incorporation of additional mechanisms. The optimal model, designated as Model 5, was obtained by further including the effect of insulin resistance induced by high peripheral insulin and the biphasic effect (positive at low levels, negative at high levels) of lipids on beta-cell secretory capacity. Although the individual data plots exhibit remarkable variability without a discernible common trend among the trajectories of the four data variables, Model 5 effectively captured the dynamics of fasting glucose, fasting insulin, Matsuda index, and beta-cell function index for individuals progressing to T2D (Fig. 5). This suggests that the model incorporates the essential physiological features of the glucose regulatory system. Furthermore, the model’s reliability was underscored by its successful cross-validation against the longitudinal prediabetes dataset.

Numerous pieces of research have investigated the molecular mechanisms underlying the progression of T2D. Ongoing debates persist on the role of insulin hypersecretion in disease evolution [33] and the significance of glucokinase in the treatment [34]. Mathematical modeling serves as an efficient and convenient approach for testing biological hypotheses, providing a quantitative framework to represent, analyze, and simulate biological processes. To scrutinize the controversial hypotheses in the diabetes field, a collection of models, each built upon different assumptions, is required for comparison and subsequent examination with sufficient data. For conciseness, we presented only one application of the model to illuminate how the model work can contribute to the study of T2D. With the credibility of our model established, we will employ it to address specific questions of interest in our future work.

## Limitations of study

In this study, we aimed at selecting the simplest model that encompasses the indispensable mechanisms to characterize the long-term T2D progression, neglecting the consideration of excessive glucagon secretion. Hyperglucagonemia can exacerbate hyperglycemia by augmenting hepatic glucose production. Recent studies have also highlighted a paradoxical effect where glucagon amplifies insulin secretion after meals, exerting a net hypoglycemic effect postprandially [35, 36]. Thus, incorporating the intricate glucagon signaling pathway into our model would inevitably increase its complexity.

Focusing on delineating the long-term dynamics of metabolic variables at their basal states, we omitted the postprandial state dynamics. Besides glucagon, postprandial glucose levels are heavily influenced by the rates of appearance in plasma of ingested glucose (Ra) and incretin effects, which are both highly variable among subjects. Consequently, extending our model to include the postprandial state would significantly increase the number of estimated parameters. While data from OGTT depict fast dynamics of the postprandial state and provide more accurate information about disease status, fitting longitudinal OGTT data simultaneously for all individuals poses a daunting computational challenge. These hurdles necessitate thorough investigation and are hence deferred to future studies.

## 6 Models and rationale

### 6.1 Starting point of model development

#### 6.1.1 Calibration for Topp’s model

The main issue with the parameter values in Topp’s model lies in the incorrect scales of G_in_, S_G_ and S_I_. The parameter G_in_ here encompasses both meal ingestion and endogenous glucose production (EGP). It is worth noting that the EGP with basal insulin is on the scale of 1060 mg/dl per day [37, 38], which is already higher than the value of G_in_ (864 mg/dl/day) in Topp’s model. When constructing a model to investigate the long-term progression of fasting glucose, it is imperative to recognize that G_in_ must include the impact of meal ingestion since the normal fasting glucose level cannot be maintained over months solely by EGP. Moreover, based on the daily energy intake and carbohydrate percentage of SWNA [39, 40], one can calculate the glucose intake from meals per day should approximate 2755 mg/dl. Therefore, G_in_ is bounded between 1060 and 3815 and can be estimated with proper values of S_G_ and S_I_.

The values of S_G_ and S_I_ given in the paper that introduced the model [41] yield a ratio that is too small (S_G_ /S_I_ = 2), whereas studies reporting that 75-85% of whole-body basal glucose uptake is by non-insulin-dependent[42] imply that the value of S_G_ /S_I_ should exceed 15. Drawing upon the findings of Tanaka et al. [43], we have adjusted the value of S_G_ and S_I_ to 17 and 1, respectively. Subsequently, G_in_ was calculated to be 2560 mg/dl/day by setting fasting glucose to 80 mg/dl in the steady state.

To better accommodate fasting glucose data with the model of Topp et al, the values for 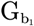 (100 mg/dl) and 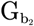 (250 mg/dl) need to be reduced. The parameter 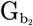 represents the critical value of glucose concentration, beyond which the beta-cell mass deteriorates due to glucotoxicity. Experimental studies indicate that the decrease in beta-cell mass begins during prediabetes [44, 45], suggesting the value of 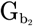 could be less than 100 mg/dl. Upon scrutinizing the complete dataset from SWNA, we observed that the fasting glucose level averaged over the population corresponding to the 2-h glucose diagnostic threshold for T2D is significantly lower than the fasting glucose threshold (partially depicted in Figure 8). As the SWNA T2D progression dataset was selected based on the 2-h glucose threshold, we chose 90 mg/dl as a reasonable value for 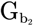 to appropriately model the fasting glucose data.

**Figure 8.**
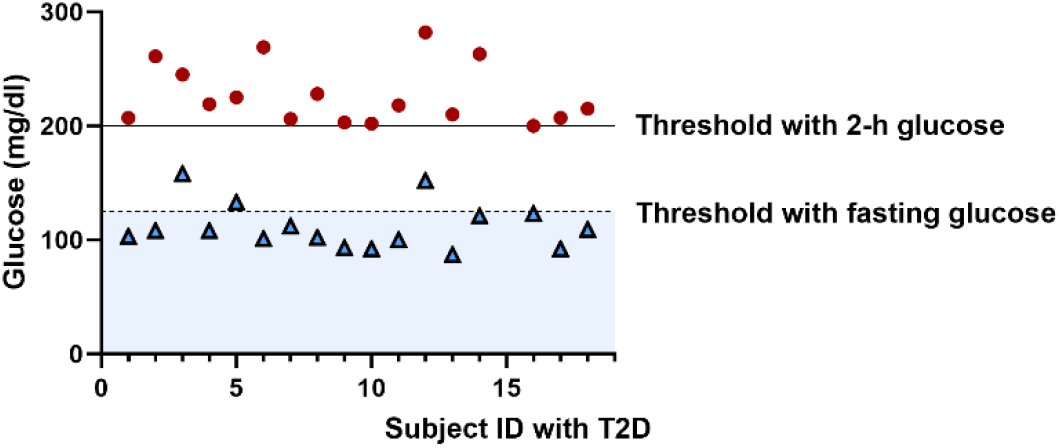
Individual fasting glucose levels recorded upon the initiation of 2-h glucose levels crossing the diagnostic threshold for T2D. Data sourced from the SWNA T2D progression dataset. The graph demonstrates that the average fasting glucose level corresponding to the 2-h glucose diagnostic threshold is significantly lower than the fasting glucose threshold within this dataset. This conclusion remains consistent for the complete SWNA dataset (data not displayed).

Lastly, the parameter values for insulin clearance rate K and time constant τ_β_ were adopted from [6]. We employ the non-linear mixed-effect (NLME) modeling approach to assess the efficacy of the Topp model using the T2D progression dataset from SWNA. The statistical models for NLME, structural model equations, and fitting results for all models discussed in this paper are available in the supplementary materials. As demonstrated in the report file for the Topp model, the individual trajectories for each variable presented in the data are poorly described by the model. We then embarked on constructing a new modeling framework to enhance performance.

#### 6.1.2 Base model construction

We initiate our model construction by changing the formulation of the insulin sensitivity function in Topp’s model. Numerous studies have shown that insulin sensitivity decreases with FFA. To quantify the individual level of excess FFA (denoted as X in the model), we first obtained the maximum and minimum PFAT values (52% and 7%, respectively) of the entire cohort across all visits. We define X to be 0 at the minimum value and 1 and the maximum value:

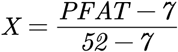

To underscore the importance of formulating insulin sensitivity as a function of PFAT rather than being solely dependent on age/time, we utilized a linear mixed-effect model to assess the relationship between Matsuda (as the response variable), PFAT, and age (as independent variables), with subject ID as the group variable and disease state as a covariate. The results (available in supplemental materials) demonstrate that both PFAT and age are significantly correlated with Matsuda, even after adjusting for glucose tolerance and repeated measures (Figure 9). Notably, the fixed effects coefficient for PFAT is four times that of the coefficient for age.

**Figure 9.**
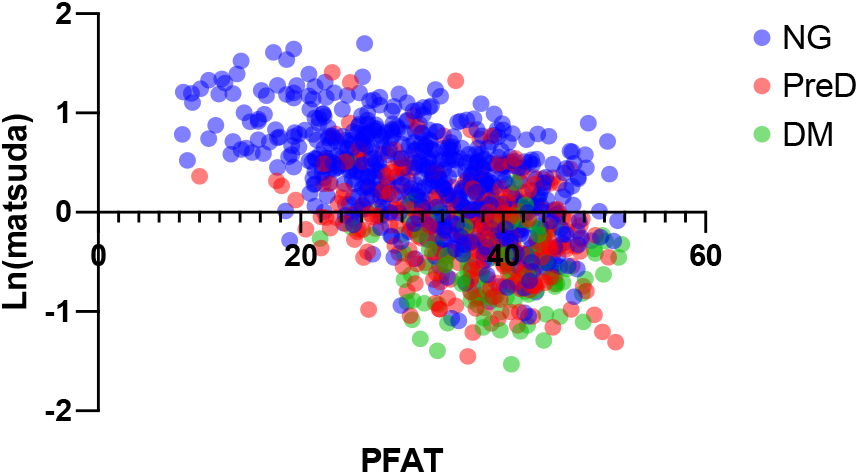
Data plot for PFAT VS log (Matsuda). The results demonstrate insulin sensitivity decreases as PFAT increases across all glucose tolerance status. The disease state was categorized by 2-h glucose levels. NG stands for normal glucose; PreD denotes prediabetes; DM stands for diabetes.

As PFAT is a dominant factor for insulin sensitivity, our initial formulation of the insulin sensitivity function depends solely on X:

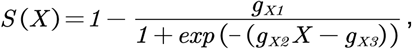

Where the parameter g_x1_ determines the extent to which the excessive FFA level can deteriorate insulin sensitivity (SI), and g_x2_ controls the steepness of the negative dependence of SI on X (Figure 10(a)). As the extent and the rate of the decline vary among individuals, the two parameters would be individually estimated using the Matsuda data.

**Figure 10.**
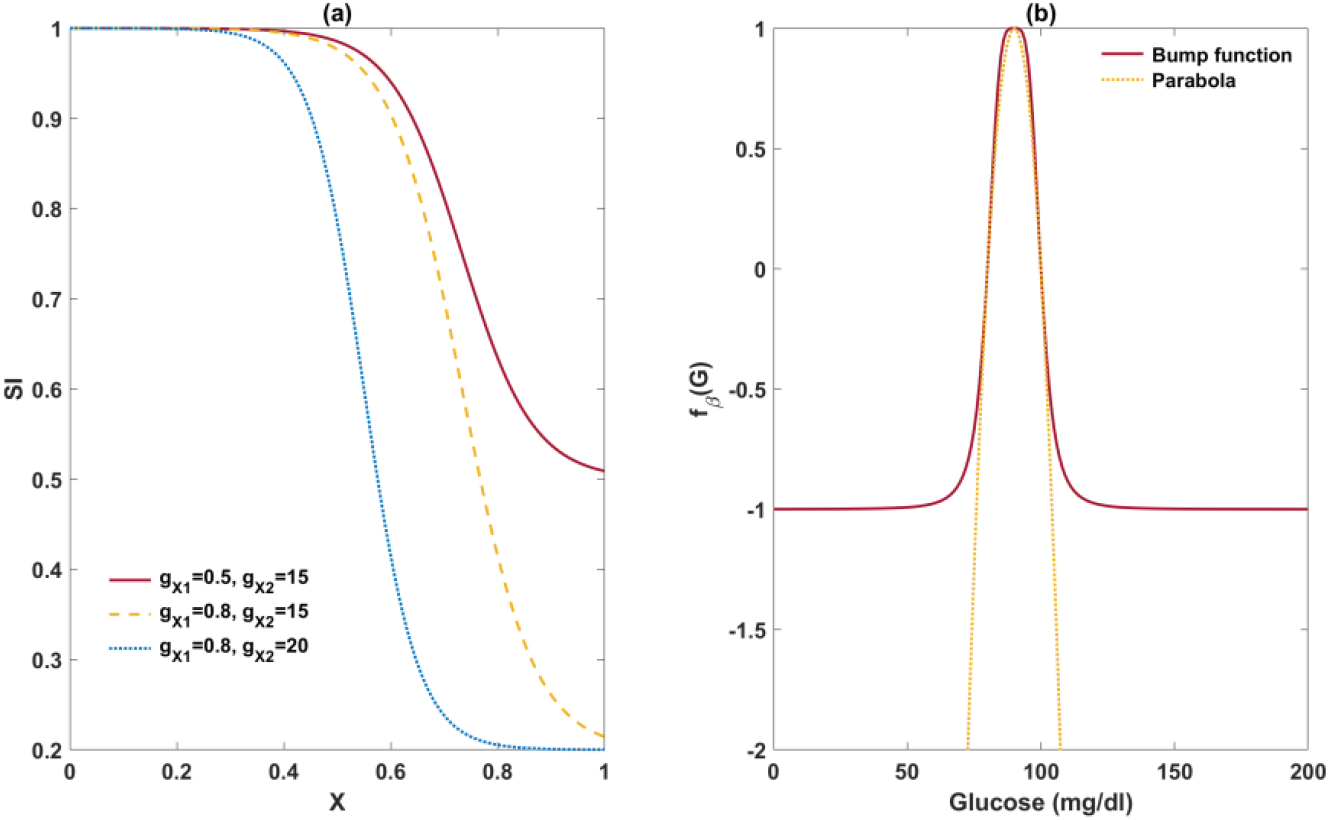
Illustrations of new functions in Model 1. Graph (a) shows the magnitude and rate of decrease of insulin sensitivity with respect to the excessive FFA level can be controlled by the parameter *g*_*X*1_ and *g*_*X*2_, respectively. Graph (b) illustrates the disparity between the beta-cell mass growth functions in Model 1 and in Topp’s model, both of which share the same roots.

In Topp’s model, the net growth rate of beta-cell mass is modeled by a parabolic function of glucose. This signifies that mild elevation in glucose can be mitigated by a compensatory increase in beta-cell mass, while high glucose levels are toxic to beta-cells, leading to a decrease in beta-cell mass. Nonetheless, the proliferative and expansion capacities of cells are finite. Considering beta-cell mass cannot expand infinitely, we replaced the exponential growth term with a logistic growth term as previously introduced by De Gaetano et al. [46]. In addition, glucotoxicity is unlikely to cause unlimited harm to beta-cell mass, so we improved the growth rate function by replacing the parabolic function with a bump function to limit the impact of glucotoxicity on beta-cell mass:

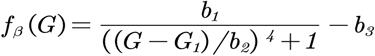

The contrast between these two functions is depicted in Figure 10(b).

With these minimal adjustments to the Topp model, we establish Model 1:

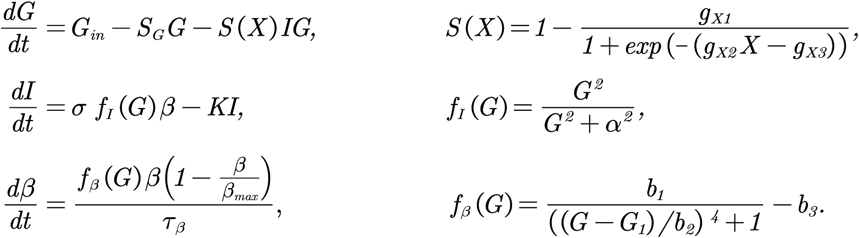

Model 1 was evaluated with the same T2D progression dataset. Although its BIC value was 31% lower than the original Topp model, the Prediction vs. Observation plots indicated that the model lacks sufficient mechanistic detail to characterize the patterns within the dataset. A model incorporating additional physiological mechanisms is desired to capture the disease progression for the subgroup of SWNA.

#### 6.2.1. Alternative model components

Within Model 1, there are eight components where modifications can be introduced to elaborate on the interactions within the glucose-insulin regulatory system. Corresponding to these components, nine enhanced functions can be applied. Detailed explanations for these adjustments are provided below.

#### 6.2.1 Incorporate the effect of aging into the insulin sensitivity function

Apart from obesity, insulin sensitivity is affected by multiple factors, including aging. The correlation analysis above revealed a significant correlation between age and the Matsuda index after adjusting for the influence of PFAT. The mechanisms of age-related insulin resistance include inflammation, oxidative stress, mitochondrial dysfunction, and changes in hormone levels and muscle mass [47-51]. Multiplying the insulin sensitivity function *S*(*X*) by a decreasing function dependent on the initial age and progression time, we obtain the function *S*(*X,AGE*):

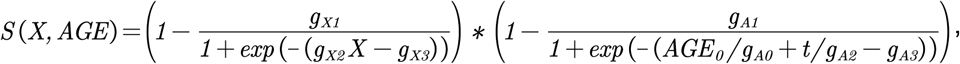

where AGE_0_ represents the age of participants at their first visit; the variable t denotes the time since the first visit; g_A3_ stands for a horizontal shift; g_A0_ and g_A2_ are scale factors; the parameter 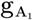 controls the magnitude of age-related insulin resistance and 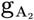 governs the rate at which insulin sensitivity decreases with age. The parameters 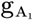 and 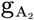 are estimated for each individual by fitting the model to longitudinal values of fasting glucose and insulin, PFAT, and the Matsuda index.

#### 6.2.2 Replace the constant glucose influx with hepatic glucose production (HGP) function

Since basal EGP is equivalent to HGP [52], and our model is designed to fit basal glucose data, these two terms are considered interchangeable in this study. In the Topp model, the rate of hepatic glucose production is assumed to be constant, which is acceptable when fasting plasma insulin is sufficiently high to compensate for the hepatic insulin resistance. However, insulin levels will inevitably decrease [53] concomitantly with an overt elevation in basal HGP as the disease progresses [27, 28]. To capture the evolution of basal HGP, we formulate the rate of HGP as a function of insulin level and excess FFAs (increase of X above 0):

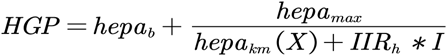

Here, IIR_h_ stands for the impact of high insulin-induced insulin resistance on hepatic insulin sensitivity and is a decreasing function of insulin, as illustrated in Figure 11(a). The influence of excess FFAs on increasing HGP, including hepatic insulin resistance and the stimulation of gluconeogenesis, is represented by a rightward shift of hepa_km_, i.e., the function hepa_km_ (X) decreases as X increases (see Suppl. Material for details). The extent to which influences HGP is determined by the parameter Hepa_SX_, which is estimated individually during the data fitting. The remaining parameter values in this function were selected to guarantee the consistency between the simulated hepatic glucose production rate and experimental data from the literature.

**Figure 11.**
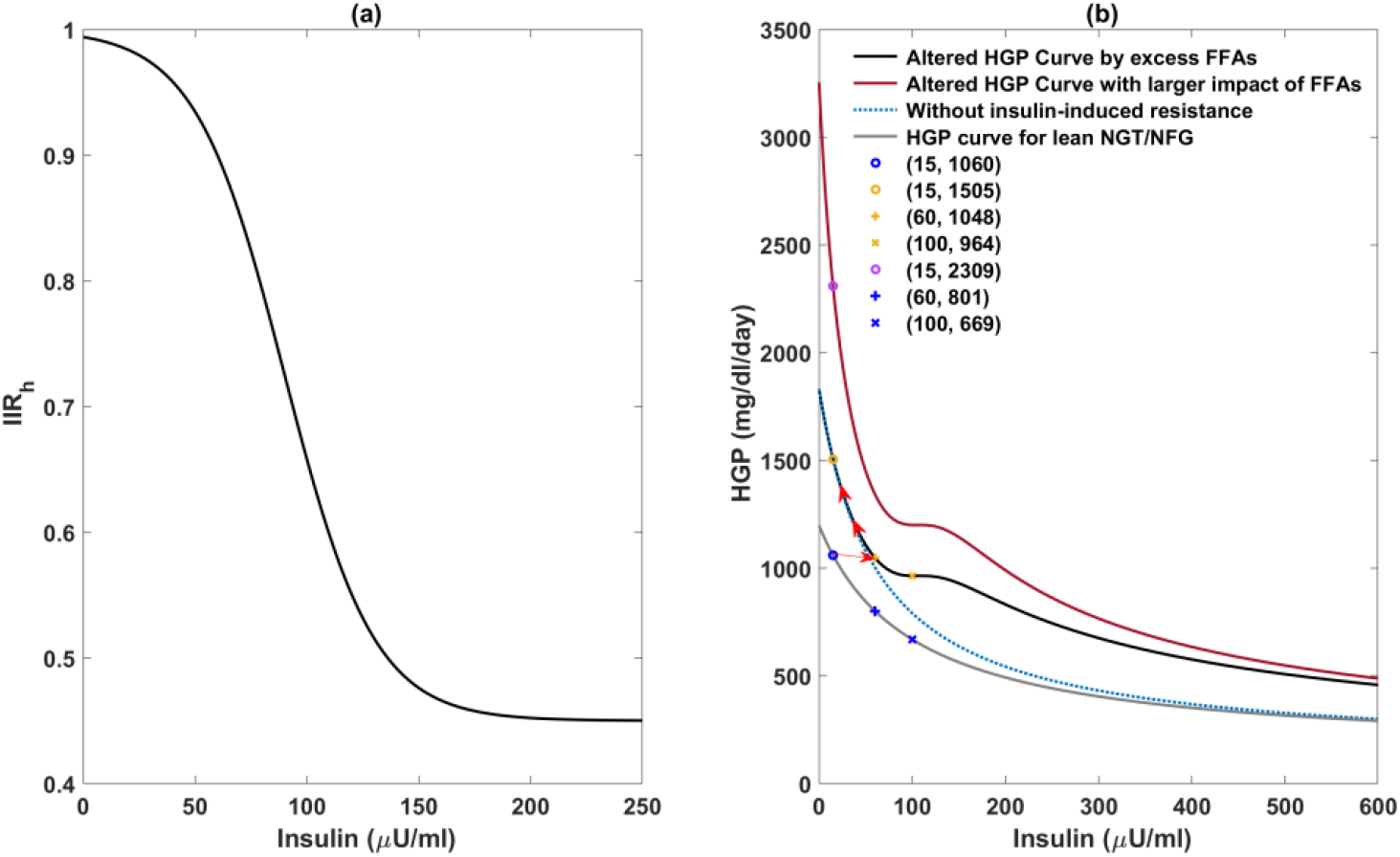
Illustration of model components related to hepatic glucose production (HGP). Panel (a) shows the effect of insulin-induced insulin resistance on HGP. The efficacy of insulin in suppressing HGP decreases as insulin increases. Panel (b) compares simulated HGP-insulin dose-response curves with and without incorporating the impact of excess FFAs (denoted by X in the model equations) or insulin-induced insulin resistance in the liver. The gray curve represents the simulated HGP-insulin dose-response curve without the influence of X or high insulin-induced resistance in the liver. The blue circle corresponds to the basal insulin and HGP for the healthy control group. The red arrow depicts the direction of basal insulin and HGP in response to the increase of X. When X is increased (higher effect of FFA, the black curve shifts further towards to red curve.

We found it necessary to incorporate X and IIR_h_ in the HGP function to ensure consistency between the HGP-insulin dose-response curve and experimental findings. In their comparison of insulin’s suppressive effect on HGP in obese T2D patients and obese control subjects, Beck-Nielsen et al. [54] proposed the observed shift of the HGP curve could be attributed to both a defect in insulin’s suppressive effect on HGP and an increase in gluconeogenesis due to elevated FFAs.

Several experimental studies have revealed that normal basal HGP can be maintained prior to the onset of diabetes, as the elevated insulin level is sufficient to offset hepatic insulin resistance [1, 29, 54]. A substantial elevation in HGP to up to twice normal occurs when insulin levels drop significantly in the diabetic stage [1, 29]. These quantitative observations can be adequately simulated by the incorporation of the hepa_km_ (X) function. Figure 11(b) shows that as X increases before the onset of diabetes, the curve shifts to the right, and the basal insulin and HGP move from the blue circle to the yellow plus initially, demonstrating normal HGP can be maintained in hyperinsulinemia. In the diabetic stage, insulin levels decline substantially, and the basal HGP moves upward along the curve. Hence, the considerable elevation of HGP can solely be observed after the onset of diabetes.

The importance of integrating IIR_h_ into the HGP function is underscored by the hyperinsulinemic-euglycemic clamp data from Basu et al [30]. They demonstrated that during the clamp, HGP can be suppressed to half the basal value for the NGT/NFG group, while the clamp HGP is only slightly lower than basal HGP for the IGT/IFG group, which has higher basal insulin levels. This reveals the efficacy of insulin to suppress HGP is impaired by elevated insulin levels, which requires incorporating IIR_h_ in the HGP function (Figure 11(b)).

#### 6.2.3 Include the effect of reduced insulin clearance

In most models examining the progression of T2D, the insulin clearance rate is assumed to remain constant for the sake of simplicity. Nevertheless, clinical findings have revealed a rapid reduction in insulin clearance when insulin secretion increases in insulin-resistance states [31]. A low insulin clearance rate has been identified as an independent risk factor predicting T2D in Hispanics, African Americans, and Native Americans [31, 55]. Thus, it is essential to incorporate the impact of reduced insulin to develop a robust model for investigating T2D progression in populations featured with high insulin secretion. Multiple clinical studies have observed an inverse relationship between insulin clearance and secretion. Additionally, data on insulin clearance rate from Mittendorfer et al. show saturation within the postprandial plasma insulin concentration range [32]. Therefore, we adopt the approach from Polidori et al., using a saturation term for insulin clearance rate:

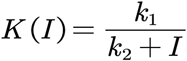

This function reflects a hyperbolic relationship between insulin clearance rates and insulin levels, aligning well with empirical data [32].

#### 6.2.4 Incorporate the high peripheral insulin-induced insulin resistance

The role of insulin resistance in driving insulin secretion is widely acknowledged. In the last two decades, mounting evidence has emerged indicating that hyperinsulinemia can in turn aggravate insulin resistance, resulting in higher insulin secretion [56, 57], at least until beta-cell failure occurs. This phenomenon is expected to be particularly prominent in the SWNA population, where euglycemic individuals with obesity exhibit hyperinsulinemia alongside severe insulin resistance. Thus, integrating the high peripheral insulin-induced insulin resistance into the insulin sensitivity function could enhance the model’s capability of capturing the insulin dynamics of SWNA. It also establishes a basis for future explorations of the contribution of insulin-induced resistance to T2D risk. To achieve this, we add an insulin-dependent component to the insulin sensitivity function with the following formulation similar to [6]:

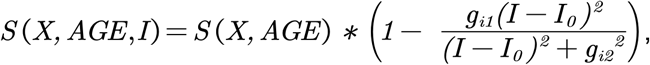

where the parameter I_0_ denotes basal insulin level; g_i1_ controls the magnitude of high peripheral insulin-induced insulin resistance.

#### 6.2.5 Integrating the influence of glucotoxicity on beta-cell secretory capacity per cell

The mechanisms underlying glucose’s stimulatory effect on insulin secretion have been extensively investigated, involving triggering and amplifying signals in beta-cells [58]. Glucose metabolism initiates the triggering signals through a series of sequential events: an increase in the ATP-to-ADP ratio, closure of K_ATP_ channels, elevation of cytoplasmic Ca^2+^ concentration, and of insulin granule exocytosis. Once glucose concentration reaches a threshold level, the efficacy of Ca^2+^ in stimulating insulin release is amplified, resulting in increased insulin secretion.

We modified the insulin equation to have Ca^2+^ concentration mediate the effect of glucose on the triggering. The stimulatory effect of Ca^2+^ on insulin release is depicted by an increasing, saturating function of glucose (f_1_ (Ca) below). The parameters in f_1_ (Ca) are estimated using data on the insulin secretion rate by glucose from human islets in vitro [59]. The corresponding fitting plot is displayed in Figure S1.

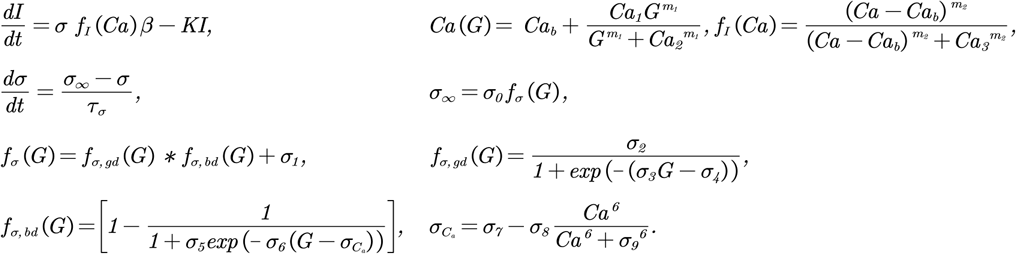

The amplifying signal is modeled as an increase in the maximum rate of insulin secretion, corresponding to the variable σ as in [6]. In the Topp model, σ was assumed to be constant, and the slow compensation for insulin resistance was assumed to occur only through an increase in beta-cell mass. However, as stated in [20] and measured in [60], even in rodents, expansion of mass alone is insufficient to explain the observed rise in insulin concentrations. Therefore, it is desirable for the model to incorporate a dynamic σ, with a rate of change that lies between fast glucose changes and slower alterations in beta-cell mass. To achieve this goal, we formulate a differential equation for σ, wherein approaches its steady-state level σ _∞_, which depends on both glucose and Ca^2+^, with a time rate 1/τ_σ_.

Although glucose acutely stimulates insulin secretion by beta-cells and chronically increases betacell mass when moderately elevated, persistent hyperglycemia is toxic to beta-cells, termed glucotoxicity [61]. Impairment of first-phase insulin secretion occurs even when glucose levels are minimally raised and become aggravated as glucose levels escalate [62]. One possible mechanism underlying glucotoxicity is via “Ca^2^_+_ toxicity”, wherein persistent inflow of Ca^2^_+_ can trigger beta-cell apoptosis [63]. Thus, we compose σ _∞_ with both a positive effect of low glucose f_σ.bd_ (G) and a harmful effect of high glucose f_σ.bd_ (G). The parameter σ_6_ modulates the sensitivity of σ to the toxicity directly from glucose, while σ_6_ represents calcium toxicity. In this work, we utilize σ_6_ to estimate individual susceptibility to glucotoxicity. The influence of σ_6_ on σ _∞_is illustrated in Figure 12(a).

**Figure 12.**
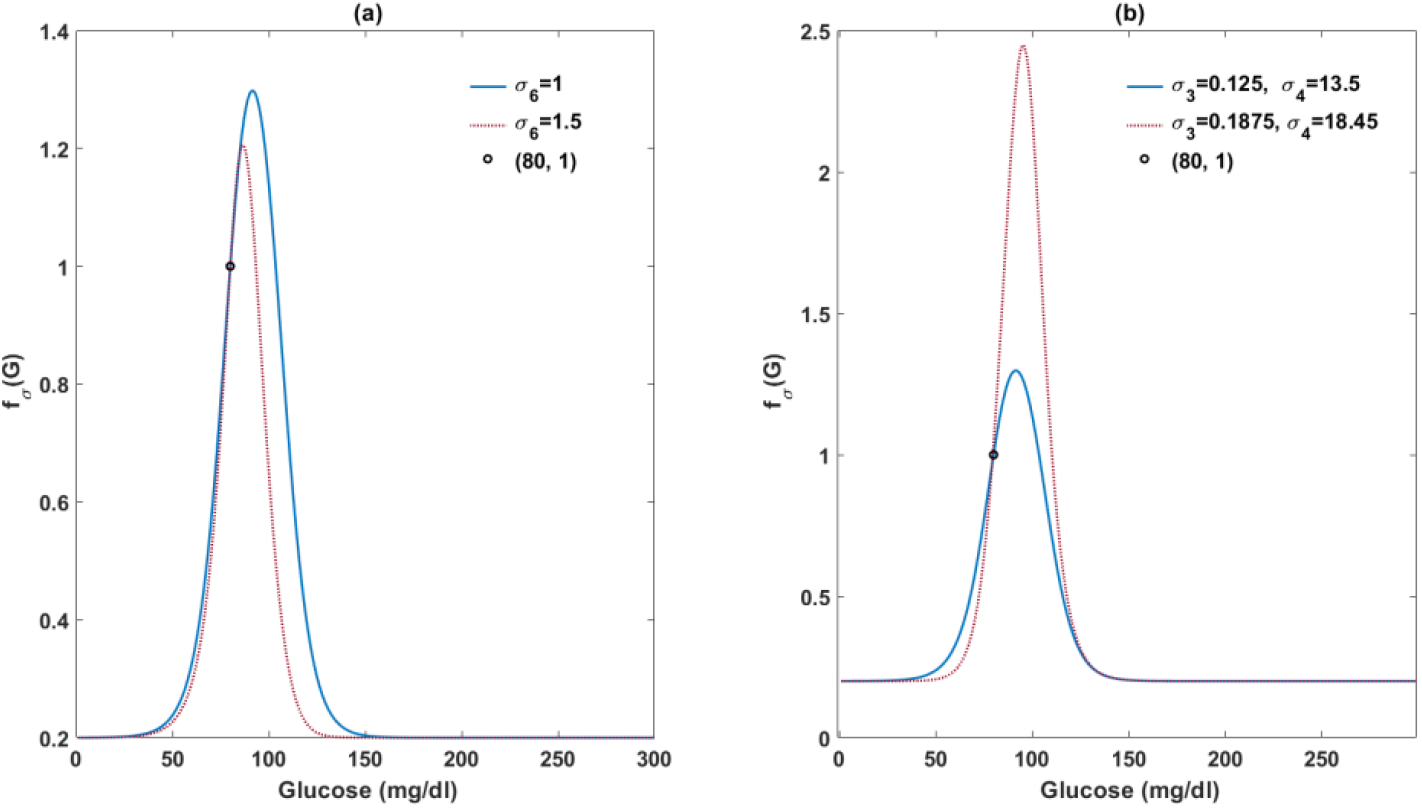
Illustration of varied beta-cell response to glucose with different parameter values in *f*_σ_(*G*). In panel (a), the circle symbolizes the default beta-cell response to basal glucose levels. A mild rise of glucose from the basal level can amplify beta-cell response. Whereas beta-cell function deteriorates as glucose levels exceed a certain threshold. An increase in σ_6_ can lead to an early occurrence of glucotoxicity. Panel (b) shows that an elevation in both σ_3_ and σ_4_ results in an augmented positive effect of glucose on beta-cell secretory capacity.

#### 6.2.6 Enhancing the positive effect of glucose on beta-cell secretory capacity

The SWNA population is characterized by high levels of insulin secretion. To explore whether this high secretory capacity can be explained by increased sensitivity to glucose stimulation in beta-cells, we raised the parameter values of σ_3_ and σ_4_ (Figure 12(b)), and then refitted the model to the data to assess whether the model performance improves.

#### 6.2.7 Including the effect of lipotoxicity on beta-cell mass

In addition to glucose, insulin secretion is impaired by exposure to FFAs, a phenomenon termed lipotoxicity. Numerous experimental studies have delved into the mechanisms through which sustained excessive FFAs impair beta-cell function[64]. Lipotoxicity can also impair compensatory beta-cell proliferation, diminishing beta-cell mass [65]. Therefore, we separate the lipotoxic effects on secretory capacity and mass. The latter is represented by a negative term, f_β,2_ in the beta-cell growth rate function:

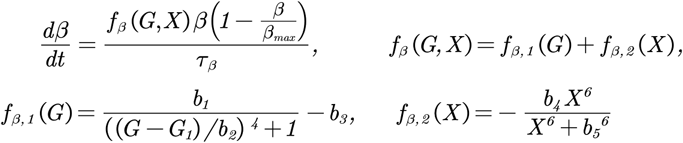

#### 6.2.8 Including merely the impact of lipotoxicity on beta-cell secretory capacity

As the prevailing evidence supports mainly the lipotoxic effect on beta-cell secretory capacity rather than beta-cell mass, we omitted the influence of lipotoxicity on beta-cell mass, thereby reducing the number of estimated parameters. The effect on secretory capacity is represented by f_σ,2_(X) in the σ _∞_ function, whereby σ _∞_ decreases with X:

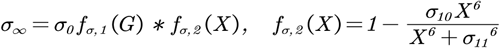

#### 6.2.9 Incorporating the lipid biphasic-effect on beta-cell secretory capacity

There is also evidence that FFA can exert a positive influence on beta cell function, contingent upon factors such as concentration, duration, and glucose levels [8, 22]. We express the biphasic effect of lipids on function by modifying f_σ,2_(X) as follows:

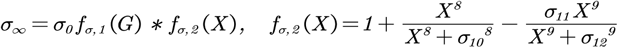

The parameter σ_11_ determines the threshold value of X that triggers lipotoxicity, as illustrated in Figure 13 (a). Figure 13 (b) depicts the balanced effects of FFAs and glucose on beta-cell secretory capacity. In the presence of high glucose, a substantial increase in X further diminishes σ_∞_, representing a synergistic impact of glucose and FFAs in causing toxicity.

**Figure 13.**
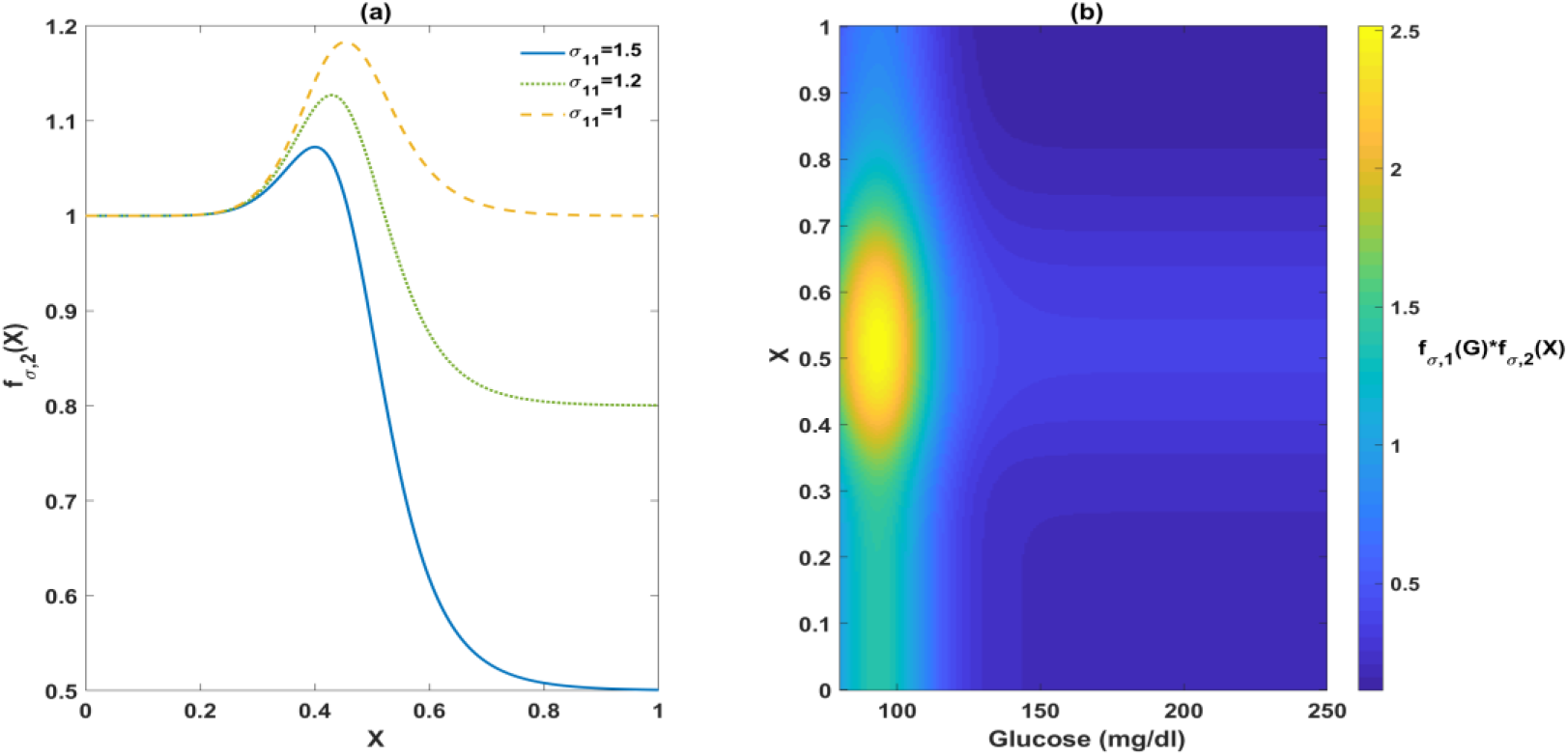
Illustration of the biphasic effect of FFAs on σ_∞_. Panel (a) demonstrates that the impact of FFAs on σ_∞_ can be adjusted by changing the parameter σ_11_. Panel (b) illustrates the balanced effects of FFAs and glucose on σ_∞_. The region where *f*_σ,1_(*G*) * *f*_σ,2_(*X*) is larger than 1 represents a positive net effect of FFAs and glucose on insulin secretion. The yellow region highlights the pair of glucose and X that generate the strongest stimulatory effect.

## Acknowledgements

The data from Southwestern Native Americans was kindly provided by Dr. Clifton Bogardus (NIDDK). The authors would like to thank the community members who participated in the study. We would also like to thank the Tribal Research Review Committee and Social Standing Committee for their review and input into the manuscript.

## Author Contributions

Conceptualization: [Boya Yang], [Arthur Sherman]; Methodology: [Boya Yang], [Arthur Sherman]; Formal analysis and investigation: [Boya Yang], [Arthur Sherman]; Writing - original draft preparation: [Boya Yang], [Arthur Sherman]; Writing - review and editing: [Boya Yang], [Arthur Sherman]; Funding acquisition: [Arthur Sherman]; Resources: [Boya Yang], [Arthur Sherman]; Supervision: [Arthur Sherman].

## Resource Table

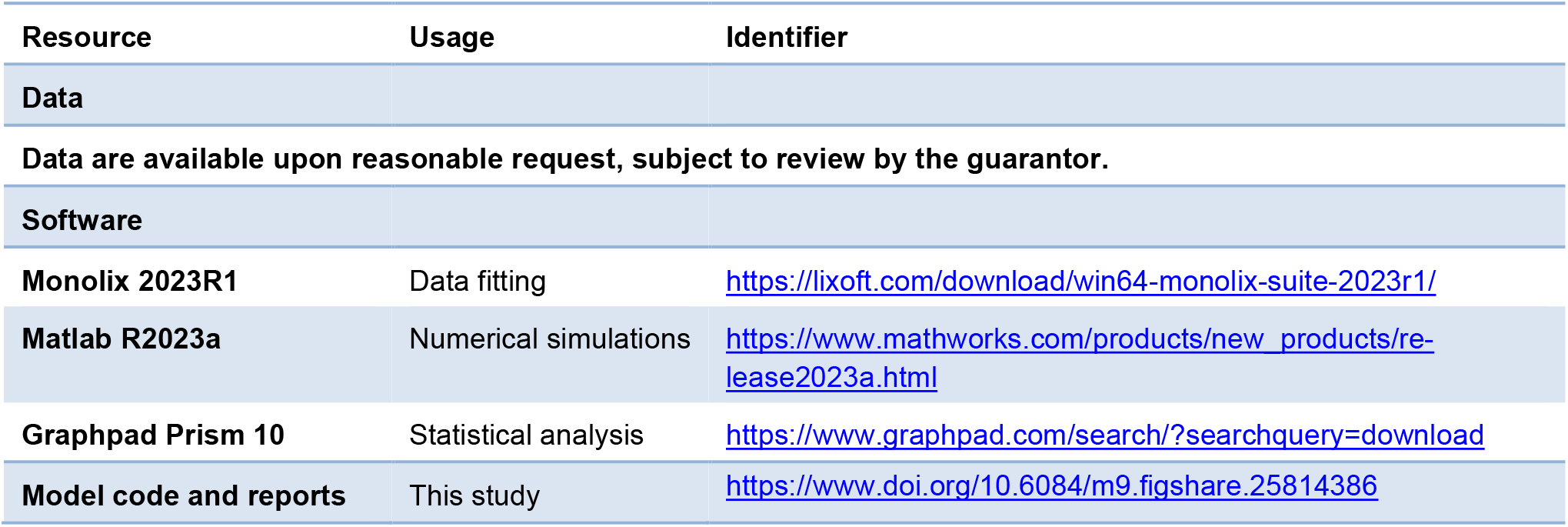

